# BROADLY NEUTRALIZING ANTI- HIV-1 ANTIBODIES DO NOT INHIBIT HIV-1 ENV-MEDIATED CELL-CELL FUSION

**DOI:** 10.1101/2021.06.08.447628

**Authors:** Nejat Düzgüneş, Michael Yee, Deborah Chau

**Author notes:** Address for correspondence: Department of Biomedical Sciences, Arthur A. Dugoni School of Dentistry, University of the Pacific, 155 Fifth Street, San Francisco, CA 94103, USA.

## Abstract

PG9, PG16, PGT121, and PGT145 antibodies were identified from culture media of activated memory B-cells of an infected donor and shown to neutralize many HIV-1 strains. Since HIV-1 spreads via both free virions and cell-cell fusion, we examined the effect of the antibodies on HIV-1 Env-mediated cell-cell fusion. Clone69TRevEnv cells that express Env in the absence of tetracycline were labeled with Calcein-AM Green, and incubated with CD4^+^ SupT1 cells labeled with CellTrace™ Calcein Red-Orange, with or without antibodies. Monoclonal antibodies PG9, PG16, 2G12, PGT121, and PGT145 (at up to 50 μg/mL) had little or no inhibitory effect on fusion between HIV-Env and SupT1 cells. By contrast, *Hippeastrum hybrid* agglutinin completely inhibited fusion. Our results indicate that transmission of the virus or viral genetic material would not be inhibited by these broadly neutralizing antibodies. Thus, antibodies generated by HIV-1 vaccines should be screened for their inhibitory effect on Env-mediated cell-cell fusion.

## INTRODUCTION

Human immunodeficiency virus type 1 (HIV-1) infection of CD4^+^ cells occurs via both free virions and cell-cell transmission. The specific interaction of the HIV-1 envelope protein, Env (gp120/gp41), with the CD4 molecule on host cells is required for both mechanisms of infection. Entry of HIV-1 into target cells requires formation of a complex between the viral SU protein, gp120, the primary receptor CD4 on the target cells, and a co-receptor. The two chemokine receptors CXCR4 (X4) and CCR5 (R5) have been identified as the principal coreceptors for X4 (T-cell-tropic virus; “T-tropic”) and R5 (macrophage-tropic virus; “M-tropic”) viruses, respectively. Binding of gp120 to the N-terminal domain of CD4 induces a pH-independent conformational change in both molecules, resulting in the interaction of the fusion domain at the N terminus of the TM protein, gp41, with the target cell membrane, and the fusion of the viral and target cell membranes.

A number of broadly neutralizing antibodies (bnAbs) were identified from culture media of activated memory B cells of an infected African donor, and were shown to neutralize a large variety of HIV strains (Walker et al., 2009) (Table 1). They recognize conserved epitopes on the variable V2 and V3 loops of the viral envelope protein, gp120. Among these bnAbs, PGT121 is one of the most potent, and protected rhesus macaques against a high dose of chimeric simian-human immunodeficiency virus (Moldt et al., 2012). A serum concentration of PGT121 in the single-digit μg/mL could mediate sterilizing immunity against the virus.

**Table 1.** Neutralization activity of antibodies against HIV-1 from different clades^#^

|  | Median IC<sub>50</sub> (µg/ml) against viruses neutralized with an<br>IC<sub>50</sub> < 50 µg/ml |  |  |
| --- | --- | --- | --- |
| Clade | 2G12 | PG9 | PG16 |
| A | 17.10 | 0.16 | 0.11 |
| B | 0.82 | 0.43 | 0.70 |
| C | 2.93 | 0.22 | 0.25 |
| D | 7.71 | 0.10 | 0.02 |
| G | 31.03 | 0.29 | 1.21 |
| F | 9.23 | 0.09 | 0.08 |

<sup>#</sup> Between 15 and 31 viruses were tested for each of the clades (Data from Walker et al., 2009).

HIV-1 can infect host cells as a cell-free virion or via cell-cell contact. HIV-1 follows different routes of cell-to-cell transmission (Dufloo et al., 2018). One mechanism involves a structure termed the “virological synapse,” whose organization requires both cellular and viral proteins. Other pathways include nanotubes, filopodia, phagocytosis or endocytosis. Cell-to-cell transmission of HIV-1 may facilitate evasion of the immune system and contribute to viral spread. bnAbs may bind conserved regions of Env, including the CD4-binding site, the glycans of the gp120 V1/V2 and V3 loops, the gp41 membrane proximal external region, and the region of interaction between gp120 and gp41.

Here, we examined whether the bnAbs PG9, PG16, 2G12m PGT121 and PGT145 inhibit HIV Env-mediated cell-cell fusion in a model system involving adherent Env-expressing HeLa cells and highly CD4+ SupT1 cells in suspension.

## MATERIALS AND METHODS

### Cells

Clone69TRevEnv cells were obtained from the AIDS Research and Reference Reagent Program, Division of AIDS, NIAID, NIH (ARRRP; Germantown, MD), and had been donated by Dr. Joseph Dougherty. The cells were cultured at 37°C, 5% CO_2_ in Dulbecco’s modified Eagle’s MEM, supplemented with 10% (v/v) heat-inactivated fetal bovine serum (FBS), L-glutamine (4 mM) (all from the UCSF Cell Culture Facility, San Francisco, CA), geneticin (200 μg/ml), hygromycin B (100 μg/ml; GIBCO, Grand Island, NY) and tetracycline (2.0 μg/ml; Sigma, St. Louis, MO)(designated as “DME/10 +tet”). Env expression was induced by transferring the “Env–“cells to medium without tetracycline (“DME/10 –tet”) 7 days before the day of the experiment (Yu et al., 1996; Yee et al., 2011). The cells were then plated in 48-well plates at 2.0 × 10^5^ cells/ml and incubated for 24 h. A fusion protein, tTa, consisting of the tetracycline repressor and activation domain of the herpes simplex virus VP16 protein is induced to undergo a conformational change in the presence of tetracycline. This prevents the binding of tTa to the inducible promoter, inhibiting the expression of Rev and Env. When tetracycline is removed, Rev is expressed and facilitates the transfer of late protein transcripts to the cytoplasm, allowing Env expression on the surface of the cells (“Env+” cells) (Yu et al., 1996; Yee et al., 2011). Env can mediate membrane fusion with HeLaT4 cells, and is thus a T-tropic protein.

CD4+ SupT1 cells (Smith et al., 1984) were used as the target cells for Env-expressing cells, and were obtained from the ARRRP; these cells had been donated to the Program by Dr. James Hoxie. SupT1 cells express high levels of the chemokine receptor CXCR4, which is a co-receptor for T-tropic HIV. The observation that Env+ cells can fuse with SupT1 cells indicates that the Env protein expressed by Clone69TRevEnv cells is T-tropic (Yee et al., 2011). The SupT1 cells were maintained in a humidified 37°C incubator with 5% CO_2_ in RPMI 1640 medium supplemented with 10% FBS, penicillin (100 units/ml), streptomycin (100 μg/ml) and L-glutamine (2 mM) (RPMI/10) (all from the UCSF Cell Culture Facility).

### Fluorescence Labeling

The Clone69TRevEnv cells were labeled with Calcein AM Green (Invitrogen, Carlsbad, CA) at concentrations of 1 or 2 μM for 30 min at 37°C. The cells were washed twice with PBS to remove excess dye. SupT1 cells were labeled with CellTrace™ Calcein red-orange (Invitrogen) at concentrations of 2 or 4 μM for 30 min at 37°C. The cells were washed twice with PBS to remove the excess dye.

### The Fusion Assay

Fluorescently labeled Clone69TRevEnv cells (2 × 10^5^ cells/well in 48-well plates) were incubated for 30 min in PBS, with or without antibodies or lectins. SupT1 cells that are gown in suspension were added at a cell density of 2.0 × 10^5^/ml to wells containing the adherent HIV-Env cells incubated previously with or without tetracycline. The two types of cells were incubated for 3 h at 37°C, under 5% CO_2_. After this co-incubation, The cells were then washed twice with PBS, and observed with a 20x objective in a Nikon Diaphot inverted fluorescence microscope equipped with a dual fluorescence filter cube (Chroma, Bellows Falls, VT). Micrographs were obtained either with a QImaging camera, using the Qcapture program on an Apple Power Macintosh G3 computer, or a Jenoptik ProgRes digital camera system using ProgRes MAC CapturePro software and an Apple iMac computer. The dose-response curves were generated from duplicate wells. Other experiments were performed with triplicate wells for each condition; the results presented are representative of at least two independent experiments.

## RESULTS

### Expression of Env mediates membrane fusion and syncytium formation

HIV-Env cells that were not induced to express the HIV-1 envelope protein, Env, did not undergo fusion with SupT1 cells (Figure 1A). HIV-Env cells induced to express Env fused with SupT1 cells expressing the CD4 receptor, resulting in the generation of orange syncytia as the red and green dyes intermixed (Figure 1B).

**Figure 1.**
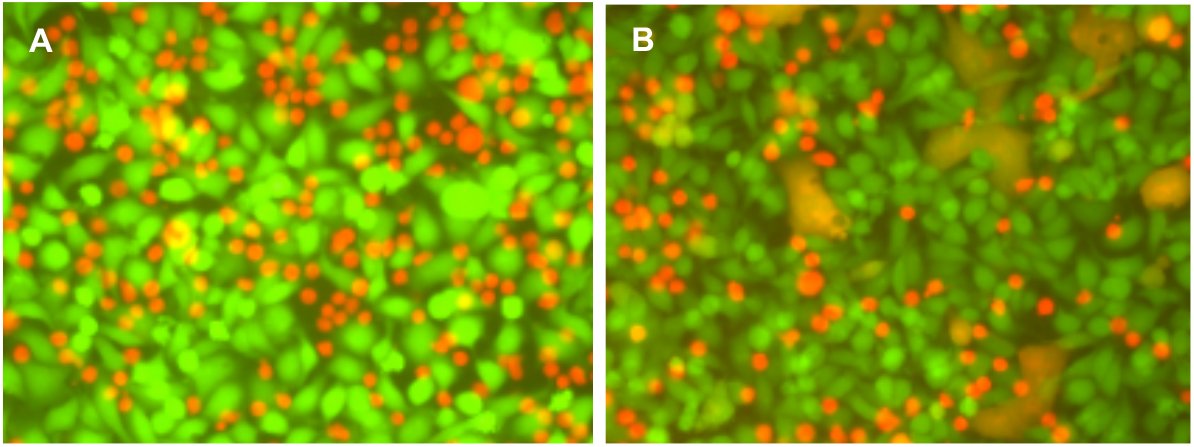
HIV-1 Env-mediated membrane fusion. **A.** HIV-Env cells *not* expressing Env (green) + SupT1 cells (red). **B.** HIV-Env cells expressing Env (green) + SupT1 cells (red). HIV-Env cells without envelope did not form syncytia with SupT1 cells. HIV-Env cells with envelope formed syncytia (orange) with SupT1 cells.

### Mannose-specific lectins inhibit membrane fusion

The mannose-binding lectins, *Hippeastrum* hybrid (Amaryllis) lectin and *Galanthus nivalis* lectin inhibited completely binding and fusion between HIV-Env and SupT1 cells (Figure 2).

**Figure 2.**
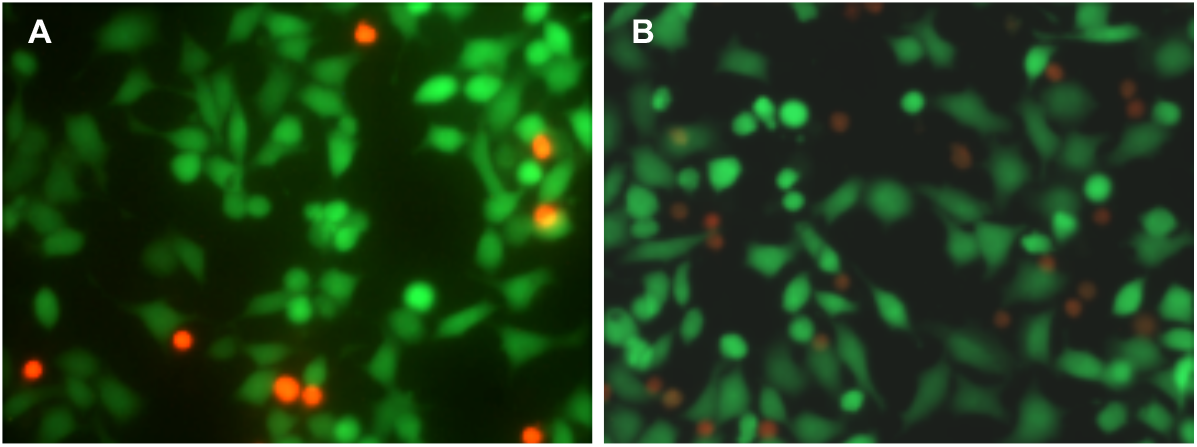
The effect of lectins on HIV-1 Env-mediated membrane fusion. **A**. *Hippeastrum* hybrid (Amaryllis) lectin (1 μg/ml) (Data from Yee et al., 2011). **B.** *Galanthus nivalis* lectin (1 μg/ml). Both lectins inhibited completely syncytia formation, as well the binding of SupT1 cells to the HIV-Env cells.

### Broadly neutralizing antibodies do not inhibit Env-mediated membrane fusion

Monoclonal antibodies PG9, PG16, and 2G12 at concentrations from 5 to 50 μg/ml had little or no inhibitory effect on fusion between HIV-Env and SupT1 cells. (Figure 3).

**Figure 3.**
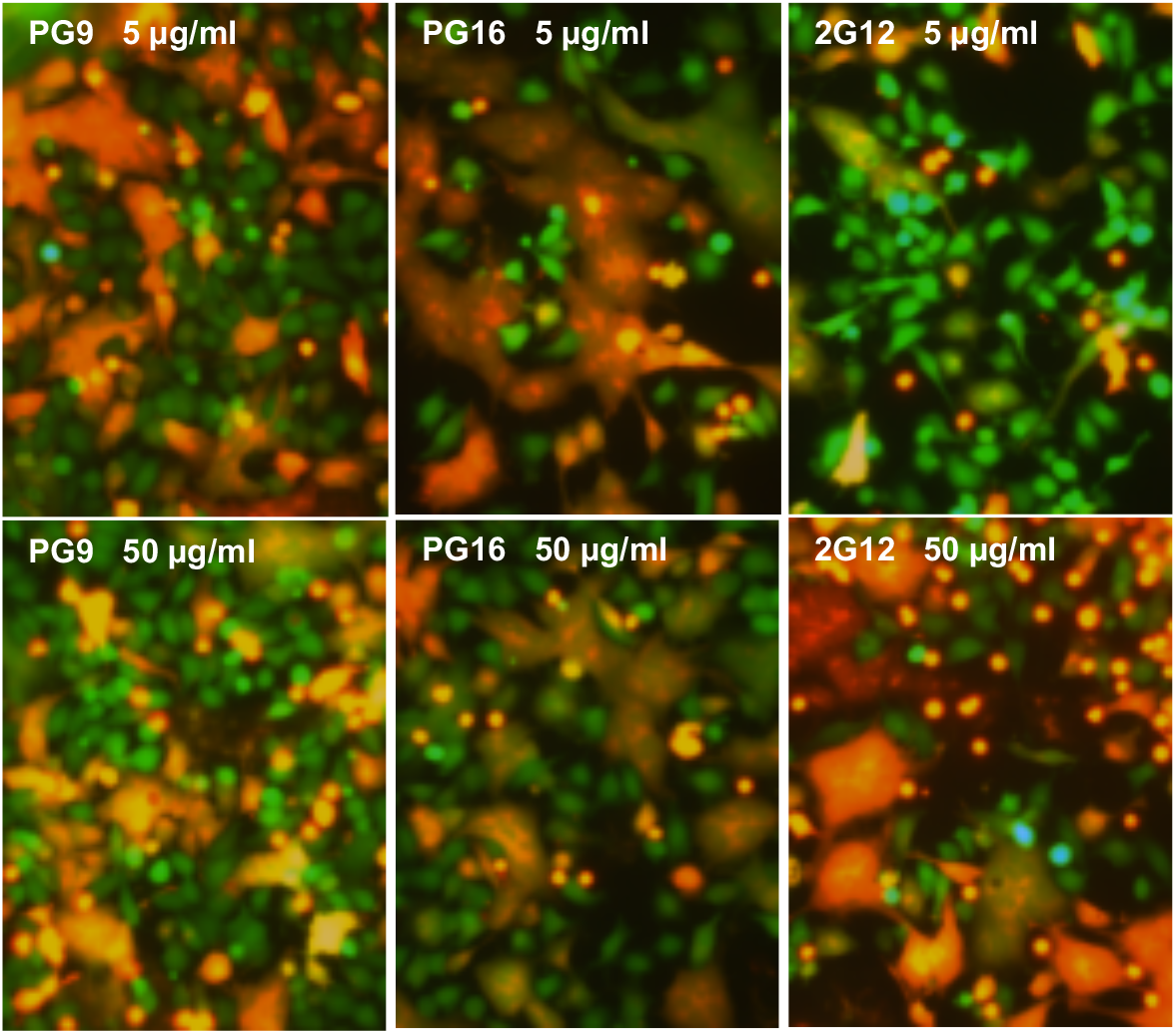
The effect of the broadly neutralizing antibodies, PG9 (left panels), PG16 (middle panels) and 2G12 (right panels) on HIV-1 Env-mediated membrane fusion. Top row, antibody concentration: 5 μg/ml. Bottom row, antibody concentration: 50 μg/ml.

Antibodies PGT121 and PGT145 appeared to have some inhibitory effect on fusion between HIV-Env and SupT1 cells at concentrations from 1 to 20 μg/ml (Figure 4). Even at 1 μg/ml PGT121, there were many red SupT1 cells that were bound to the Env+ cells that had not been able to fuse; nevertheless, some syncytia had formed. At 20 μg/ml PGT121, there were many fewer adherent SupT1 cells compared to the 1 μg/ml antibody concentration, suggesting that this antibody inhibits Env binding to its CD4 receptor on the SupT1 cells, although some syncytia were observed. PGT145 antibody was not effective in inhibiting the Env-CD4 interaction and thus did not prevent SupT1 cells from binding the Env+ cells (Figure 4).

**Figure 4.**
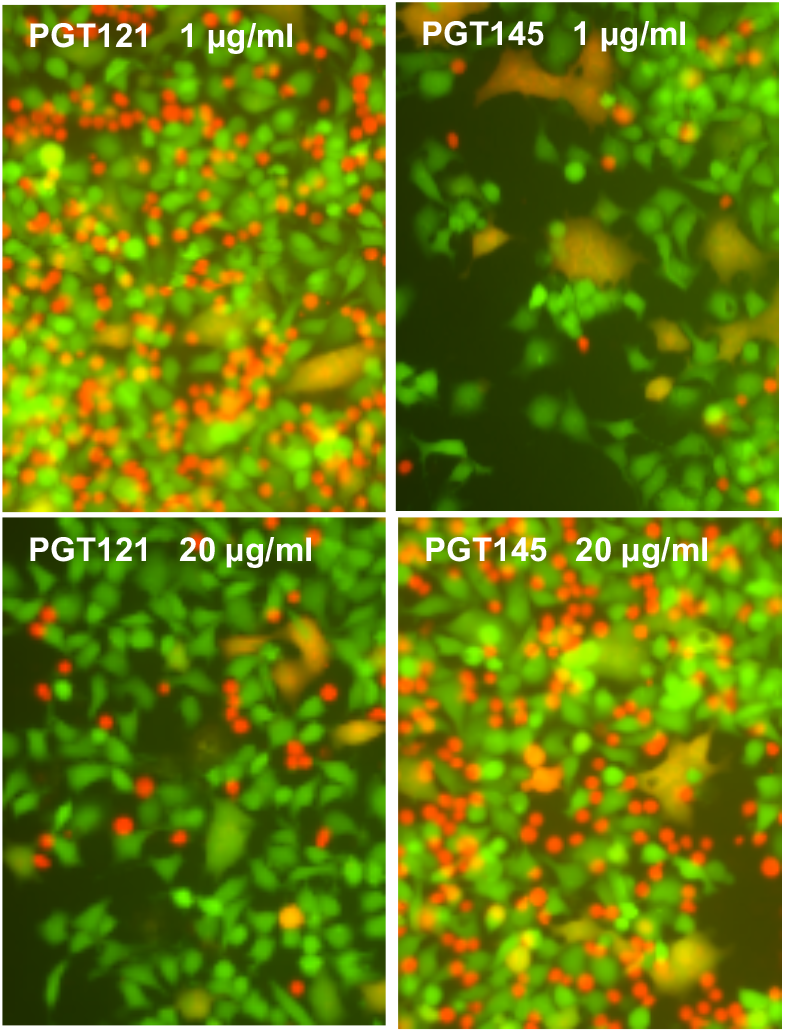
The effect of the broadly neutralizing antibodies, PGT121 (left panels) and PGT145 (right panels) on HIV-1 Env-mediated membrane fusion. Top row, antibody concentration: 1 μg/ml. Bottom row, antibody concentration: 20 μg/ml. The antibody concentrations were lower than that in Figure 3 because of the limited availability of antibodies.

## DISCUSSION

Our observation that antibodies that inhibit HIV infection are not effective against syncytium formation suggests that the mechanisms of interaction of Env with cell membrane CD4 and co-receptors may be different in cell-cell and virus-cell membrane fusion, as we have suggested previously (Konopka et al., 1995). It is also possible that the cell-cell interface is less accessible to the antibodies (Dufloo et al., 2018). Our cell-cell fusion system does not replicate the virological synapse, except for the local expression of Env alone without a budding virus, and the presence of the CD4 receptor and accompanying co-receptors.

Passive immunization with bnAb and challenge experiments in non-human primates indicate that these antibodies can protect the host against infection (van Gils & Sanders, 2013; Pegu et al., 2017). The target binding sites (epitopes) of bnAbs can be used as templates for the design of immunogens that may be able to induce similar antibodies. Genes encoding bnAbs may be delivered intramuscularly to facilitate the secretion of the antibodies (Balasz et al., 2011). Immunotherapies against HIV-1 may be used for prevention or maintenance therapy (Caskey et al., 2019).

Antibodies PG9, PG16, 2G12, PGT121, and PGT145 are ineffective against cell-cell fusion, indicating that transmission of the virus or viral genetic material from cell to cell would not be inhibited in a clinical setting. The search for immunogens to elicit antibodies that are broadly neutralizing must be reconsidered to include viral antigens that also induce antibodies that inhibit cell-cell fusion. New strides in cell-cell fusion inhibition should be toward compounds or antibodies that bind the Env protein as mannose-binding lectins do. Alternatively, the effect of bnAbs on syncytium formation between chronically infected cells and uninfected cells, or the direct transmission of HIV-1 from an infected cell to an uninfected cells would need to be investigated.

## Conflict of interest statement

The authors declare that they have no conflicts of interest.

## Notes

### Competing Interest Statement

The authors have declared no competing interest.

